# Signal processing basics applied to Ecoacoustics

**DOI:** 10.1101/2021.01.26.428328

**Authors:** Ignacio Sanchez-Gendriz

## Abstract

Ecoacoustics is a research field that has attracted attention of researchers from areas as diverse as ecology, biology, engineering, and human sciences, to cite a few. Ecoacoustics studies the sounds that emanates from the environments, by gaining insights of landscape dynamics from acoustic patterns and particularities at different places. With recent advance in technology, it is common to obtain sound datasets recorded 24-h a day for several months. The analysis of these long-term sound recordings represents several challenges. Investigators already involved in acoustic are familiar with signal processing methods, which are essential tools for sound analysis. However, in general beginners interested in Ecoacoustics do not share this background formation. The present work illustrates basic topics of digital signal processing in a comprehensible style, and an effective pipeline for long-term sound explorative data analysis (EDA) is presented. Finally, the described signal processing fundamentals are applied for EDA of 1-month underwater acoustic data recorded in a Brazilian marine protected area. The Matlab codes used for the analysis will be available as supplementary material.

## 1. Introduction

Ecoacoustics considers soundscape [1], [2] or sounds that emanates from environments as a key ecological attribute [3]. Sounds can give insights on dynamics of environments and it is recognized as an indicator of ecological processes [4]. Recent technological advance allows to collect sounds 24-h continuously for several months, which represents an opportunity for new discoveries, but also it is a challenge to overcome from the analysis point of view. The main objective of the present work is to illustrate how Digital Signal Processing (DSP) basics can be useful in the context of Ecoacoustics analysis.

Here we describe a framework that can be implemented for Explorative Data Analysis (EDA) of long-term sound recordings. The presented EDA framework is applied for 1-month underwater sound dataset.

The paper was structured as follows. Section 2 describes basic DSP concepts, exemplified by using simulated or real sound signals; at the end of the section the framework for EDA of long-term sound recordings is presented. In section 3 the application of the proposed framework is used for the analysis of 1 month of continuous underwater sound recordings. Section 4 discusses the applicability and possible extensions of the illustrated framework. The Matlab codes used for generating all the figures as well as for implementing the presented analysis will be available as supplementary material.

## 2. Methods

For beginners in areas like Ecoacoustics that deals with sound signal analysis, there are several new terms and concepts than need to be understood. A word cloud representation could be used to illustrate the feeling of overload when newcomers read technical papers for the first time, see Figure 1A. However, the concepts explained using a certain sequence and aiming to support a specific analysis could contribute to understanding and practical applications, see Figure 1B as an illustrative example.

**Figure 1.**
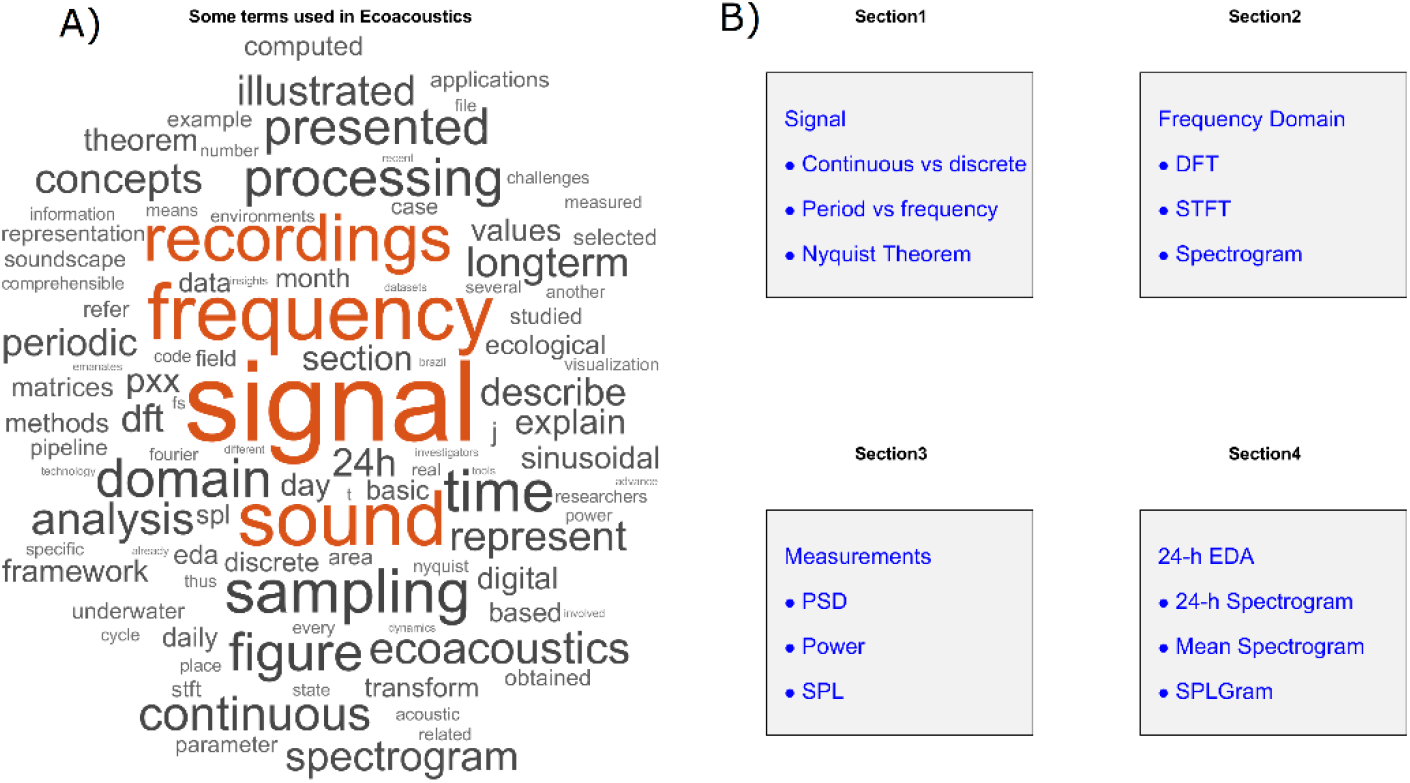
Representation of terms, concepts and topics that can be found in Ecoacoustics literature. A) Word cloud from the present manuscript. B) Some of the principal concepts covered in *Methods* section.

Methods here described are divided into five subsections, which deal respectively with signal fundamentals, frequency domain methods, filtering basics, some measurements relevant for acoustic analysis and finally the description about the proposed framework for EDA of long-term sound recordings.

### 2.1. Signal fundamentals: continuous, discrete, and sampling, period and frequency

The concept of signal as used in the present text, will refer to the sequence of values related to the magnitude of a physical variable measured by a sensor. Specifically, sound signal refers to the signal captured by a microphone (or hydrophone in case of underwater recordings). Physical variables in the nature are continuous (analog signals), they can be evaluated at every time instant, thus resulting in infinite possible values for a finite time interval measurement

Furthermore, a signal *x*(*t*) can be classified based on its periodicity as periodic or aperiodic. Periodic signals repeat its values every *T* time intervals, which can be formally written as *x*(*t*) = *x*(*t* + *T*). An example of a continuous periodic signal is illustrated in Figure 2A; for sinusoidal signals like this, the frequency of the signal (f) is the rate of cycles/sec (Hz), and the relation *T* = 1/*f* is valid.

**Figure 2.**
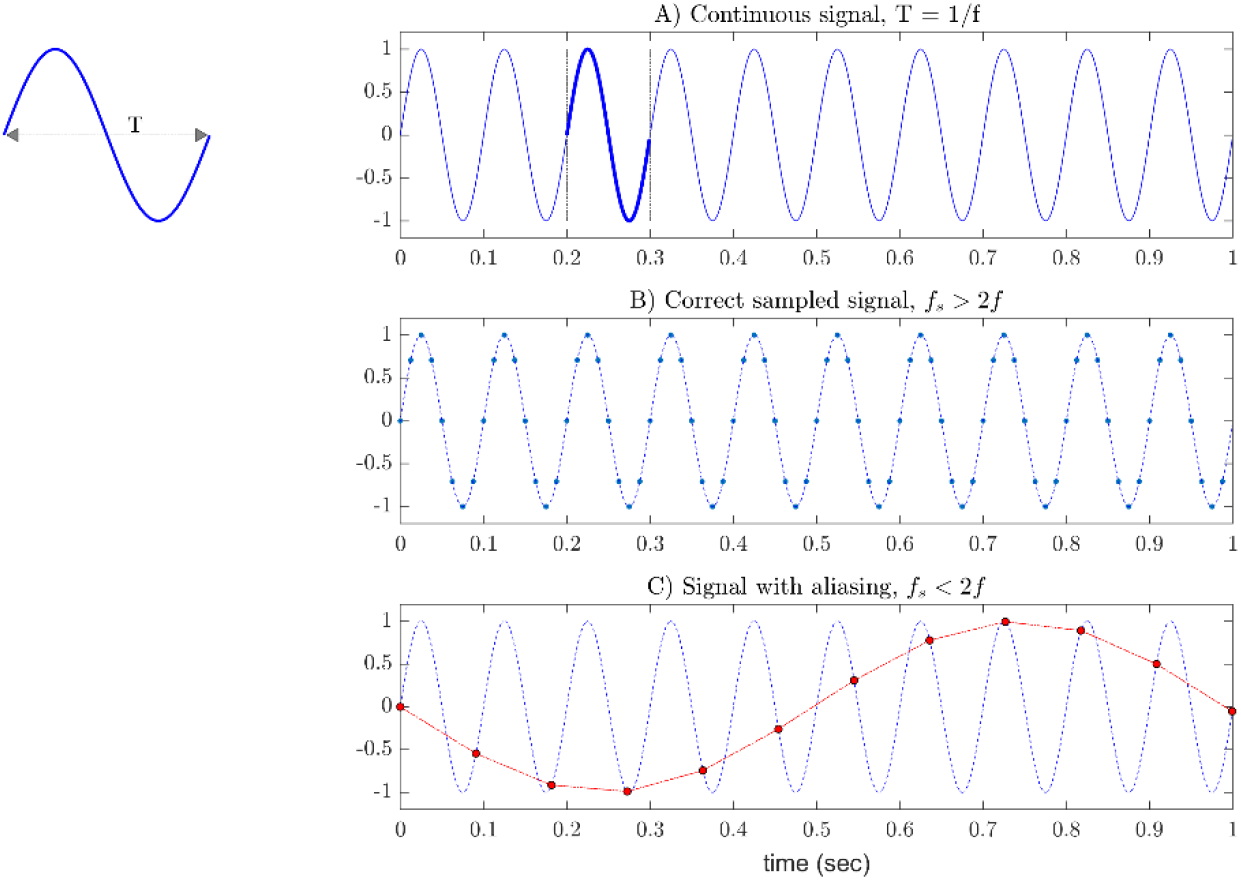
Representation of continuous and discrete periodic sinusoidal signals. A) Continuous sinusoidal signal, one cycle of the signal is highlighted in dark blue, the duration of each cycle of the signal is measured by the parameter T, which denotes the period of the signal; the number of cycles in 1 sec is the frequency (f = 10 Hz) of the signal. B) Discrete signal sampled in accordance with Nyquist’s theorem (fs > 2f), each point represents a sample from original continuous signal (dashed line). C) Discrete signal sampled not complying Nyquist’s criterium (fs < 2f).

Computers cannot manage infinite sequence of values; therefore, continuous signals must be sampled to be stored and analyzed. After the sampling process, finite precision values measured at specific time instants are obtained, i.e. a digital signal, see Figure 2B for a representation of a digital signal. The sampling process implies that not all signal values are stored, so it is correct to think that sampling could imply some loss of information, thus some precautions must be considered. The sampling frequency (*f*_*s*_) parameter defines the number of samples taken every second. The Nyquist theorem states that to perform a sampling without loss of information the sampling frequency must be selected to be superior to 2 times the highest frequency in the continuous signal being sampled [5]. Figure 2B shows a signal sampled in accordance with the sampling theorem. It can be observed from Figure 2B that the underlying (continuous) signal it is well represented by samples. In contrast, Figure 2C represent a sampling with a *f*_*s*_ below the ideal limit establish by the Nyquist theorem. For this case, the samples do not represent exclusively the frequency of the original signal, thus the Figure 2C illustrates a phenomena know as aliasing, where the frequency of the original signal is falsely presented as another frequency of lower value.

Obtaining the frequency for sinusoidal sequences it is not a complex task, however, for dealing with sound signals a more elaborated mathematical tools such as Discrete Fourier Transform (DFT) needs to be used.

### 2.2. DFT, signals from another perspective: the frequency domain

The time domain representation of a given signal illustrates how its amplitude changes over time, this representation was used in Figure 2. Based on the fact that signals can be represented as a sum of sine and cosines [6][7], if we know the amplitude and frequency of the sinusoidal ‘units’ that compose a signal, we can then represent it in another perspective: the frequency domain. Figure 3 shows the time and frequency representation for a signal x(t), obtained by the sum of five sinusoidal signals (Eq. 1).

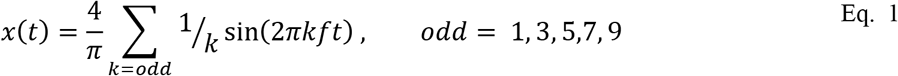

**Figure 3.**
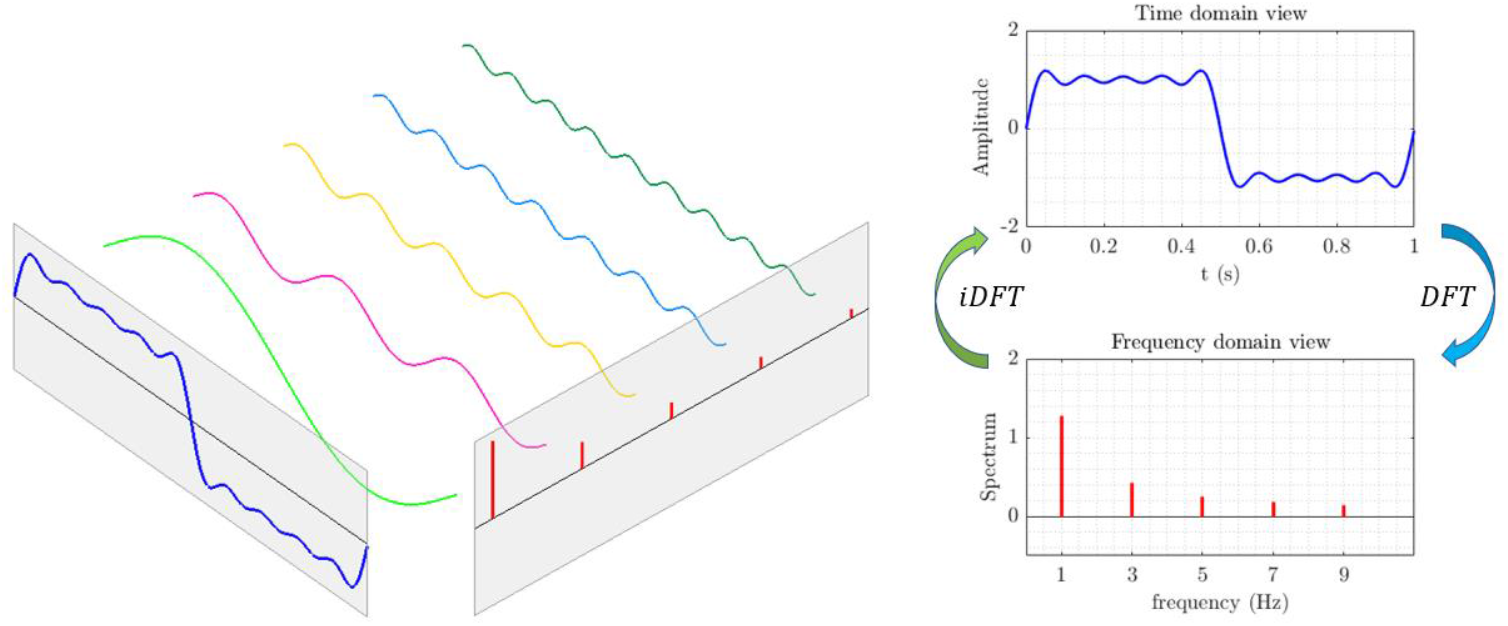
Time and frequency domain view of a signal composed by the sum of basic sinusoidal signals. DFT can be used to transform from time domain to frequency domain, in contrast inverse DFT (iDFT) allows to convert from frequency domain to time domain.

Specifically, the spectrum in frequency domain illustrates the amplitude and frequency for sinusoidals that composes a given signal. One way to transform from time domain to frequency domain and vice versa is by means of Fourier Analysis, which have been applied in ecological studies [8], [9]. Particularly for discrete signals Discrete Fourier Transform (DFT) and inverse DFT (iDFT) can be used, both DFT and IDFT are part of the Fourier Analysis methods. It is worth noting that signals in time or frequency domain contain the same information, the difference is only related with mathematical representation (just as 2+2+2 is the same as 3*2). In general terms, DFT returns complex values, the magnitude and phase of those complex numbers are defined as Magnitude and Phase spectrums, respectively. In the manuscript, the term Spectrum (without any other indication) refers to the Magnitude Spectrum, as it is the most common term used in Ecoacoustics analysis.

For a given discrete signal the *x*[*n*] with N samples, the DFT of *x*[*n*] denoted by *X*[*k*], can be written as in Eq. 2, and the iDFT formulation can be written as in Eq. 3, where *W*_*n*_ is a complex number see [6]. It can be observed that the DFT and the iDFT of discrete signals can be computed through a straightforward formulation.

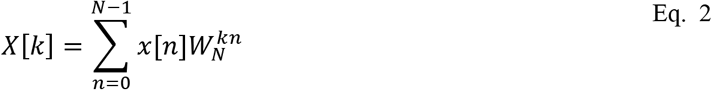

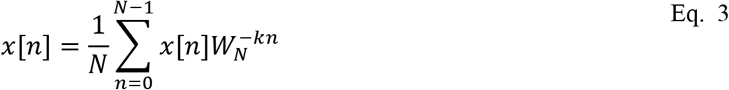

The implementation of DFT directly from its mathematical definition is computationally inefficient. For DFT to be used for practical applications, principally for massive data analysis, computational efficiency is one of the main requirements to be met. The methods designed for efficient computation of DFT are referenced in the literature as Fast Fourier Transform (FFT) algorithms [6], [10]. Thus, several software packages such as Matlab, Python and R will compute DFT by calling FFT inbuilt functions.

As we saw up to this point, DFT allows to get the frequency content for a signal. Nevertheless, when signals change frequency content over time, the DFT will not allow to distinguish the frequency variation in time. In cases that it is necessary to compute the frequency variation of signals over time, methods in time-frequency domain such as Short Time Fourier Transform (STFT) should be used [11].

#### 2.2.1. STFT and Spectrogram

The STFT of a signal is processed by sliding sections of the signal and calculating the DFT for the respective windows, thus STFT computation can be represented as an iterative process. For more details see the steps for STFT computation in [11]. Figure 4 illustrates the STFT computation process for a simulated signal (chirp) that changes its frequency over time, from 5 Hz to 15 Hz. Consecutive sliding windows allow time overlap between them and for each step the selected signal slice is multiplied by a window function. At each step, the multiplication between the sliding window and the window function produces a signal segment (windowed signal). The multiplication by the window function diminishes possible discontinuities between beginning and ending of the windowed signals, since DFT assumes that analyzed signals are periodic. This intermediary step minimizes an undesired phenomenon known as spectral leakage [6], characterized by an ‘energy leakage’ to frequencies which are not present at the original signal [12].

**Figure 4.**
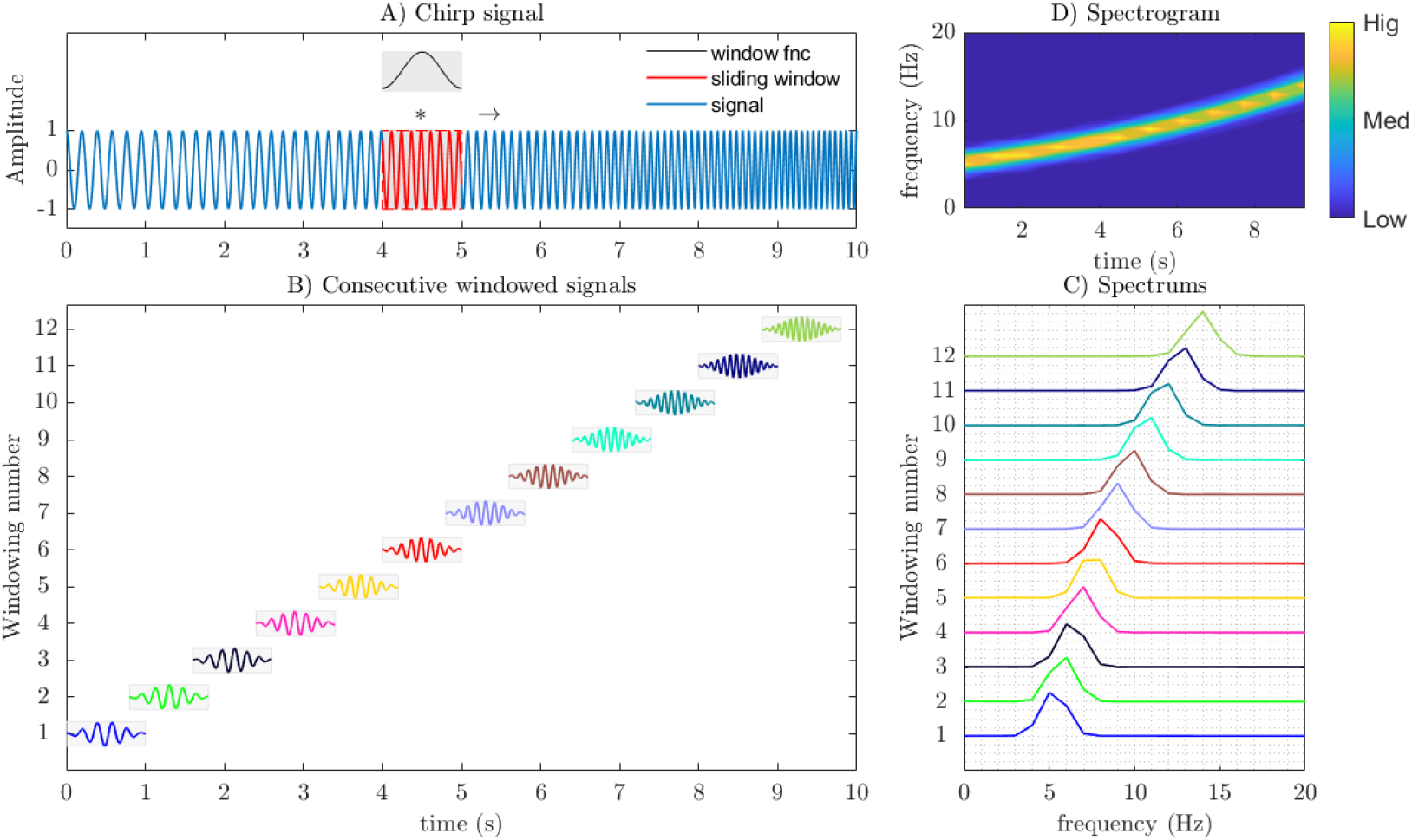
Representation of the procedure of windowing, STFT computation and Spectrogram visualization for a chirp signal with frequency varying from 5 Hz to 15 Hz. A) Nonstationary signal changing frequency content over time, a sliding window and the Hamming function are highlighted. B) Windowing procedure used to compute the STFT. C) The spectrums magnitude for each sliding window, all the computed spectrums compose the STFT for the analysed signal. D) Spectrogram for the presented signal.

For the example illustrated in Figure 4 each sliding window have 1 sec duration, consecutive segments have 20 % overlap (0.2 sec), and the Hamming^*^ function was selected as the window function. This setup generates 12 windowed signals and 12 spectrums, see Figure 4B and Figure 4C, respectively. The visualization of the amplitude of the spectrums that compose the STFT is known as the spectrogram, see Figure 4D. The spectrogram represents time along the x-axis, frequency along y-axis and amplitude of the time-frequency bins as gradient of colors, as can be observed in Figure 4D.

#### 2.2.2. Parameters and considerations about practical computation of STFT

##### Time vs frequency resolution trade off (Uncertainty principle)

When calculating DFT for each windowed signal, the frequency resolution for the respective spectrum is inversely proportional to window duration [12]. For the example shown in Figure 4B, where the length of the windows represents 1 second, the frequency resolution is 1 Hz, which means that frequency components separated by less than 1 Hz cannot be differentiated. In order to improve frequency resolution the window duration should be increased, thus decreasing time resolution. That is, there is a compromise between frequency resolution and temporal resolution. If we improve the resolution in one domain, we get worse in the other. A processing trick that can be used to improve the visualization of time resolution without a necessary detriment of frequency discrimination is by allowing overlaps between consecutive windows.

##### Visualization in dB scale

Decibel (dB) is a logarithmic scale that is commonly employed for quantifying a physical variable with large dynamic range, such is the case of sound [13]. For visualization purposes the logarithmic transformation 20log(*X*) could be employed for illustrating the spectrogram obtained from STFT computation. Also, for quantifying acoustic metrics it is common practice to represent it as a dB level relative to a reference pressure, *P*_ref_; for underwater measurements *P*_ref_ = 1*μPa*, and for terrestrial applications *P*_ref_ = 20*μPa* [14].

As we saw up to this point, signals can be characterized by its spectral content, next we will illustrate how to select desired spectral components for a given signal, while suppressing undesired frequency components. The procedure that allows such spectral separation is known as filtering.

### 2.3. Filtering basics

Filtering is an operation that produces an output signal based on the input signal. In principle, filters can be used to suppress some unwanted frequency components while maintains desired spectral content. For some researchers the filtering operation is as a black box, in which they feed a signal *x*[*n*] to the input and get a signal *y*[*n*] as the result. In this section we are going to clear up some "mysteries" of the filtering procedure’s black box. Since we are dealing with digital signals in the present manuscript, we will describe digital filters, thus when referring to filtering process we will be referring to numerical operations performed on digital samples.

#### 2.3.1. FIR and IIR filters

The Eq. 4 illustrates the filtering operation, the output at the current sample can be computed from a linear combination of current and past input samples, and past output samples. The numbers *a*_*i*_ and *b*_*j*_ are known as filter coefficients. The filter order is the maximum value between N and M, where N and M indicate the number of input and output past samples that can be used for computing the current output, respectively.

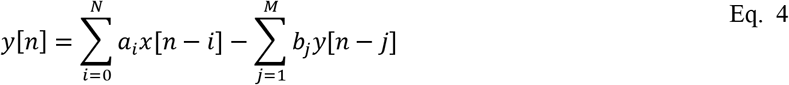

If all the *b*_*j*_ (*j* = 1,2, …, *M*) coefficients are zero, then the filter is classified as Finite Impulse Response (FIR), and the output at the current sample only depends on current and past inputs *x*[*n*]. On the other case, if at least one value of *b*_*j*_ is different from zero the filter is classified as Infinite Impulse Response (IIR), and the process include also past samples from the output.

Frequency response is one of the most important criteria for designing digital filters. Frequency response indicates how the filter select (whether it attenuates or not) specific spectral components contained in the signal x[n]. Filters can be classified based on frequency response characteristics, and the four most common categories are Low-Pass, High-Pass, Band-Pass and Band-Reject [15].

Figure 5 exemplifies a low-pass and hig-pass filtering on simulated signal composed by the sum of sinusoidals with frequencies at 10 Hz, 30 Hz, 50 Hz, 70 Hz and 90 Hz. Since filters selects which spectral components pass or are attenuated, they can be easily understood on frequency domain. When a signal is filtered, basically its spectrum is multiplied by the filter frequency response. See Figure 5 for an illustration of filtering functioning, for example, when a low-pass is applied on the signal, low-frequencies in the pass band are multiplied for values near one, thus spectral content of the pass band is practically unchanged. On the other hand, high-frequency components in the stop band are multiplied by values near to zero, therefore higher frequencies are almost eliminated after low-pass filtering.

**Figure 5.**
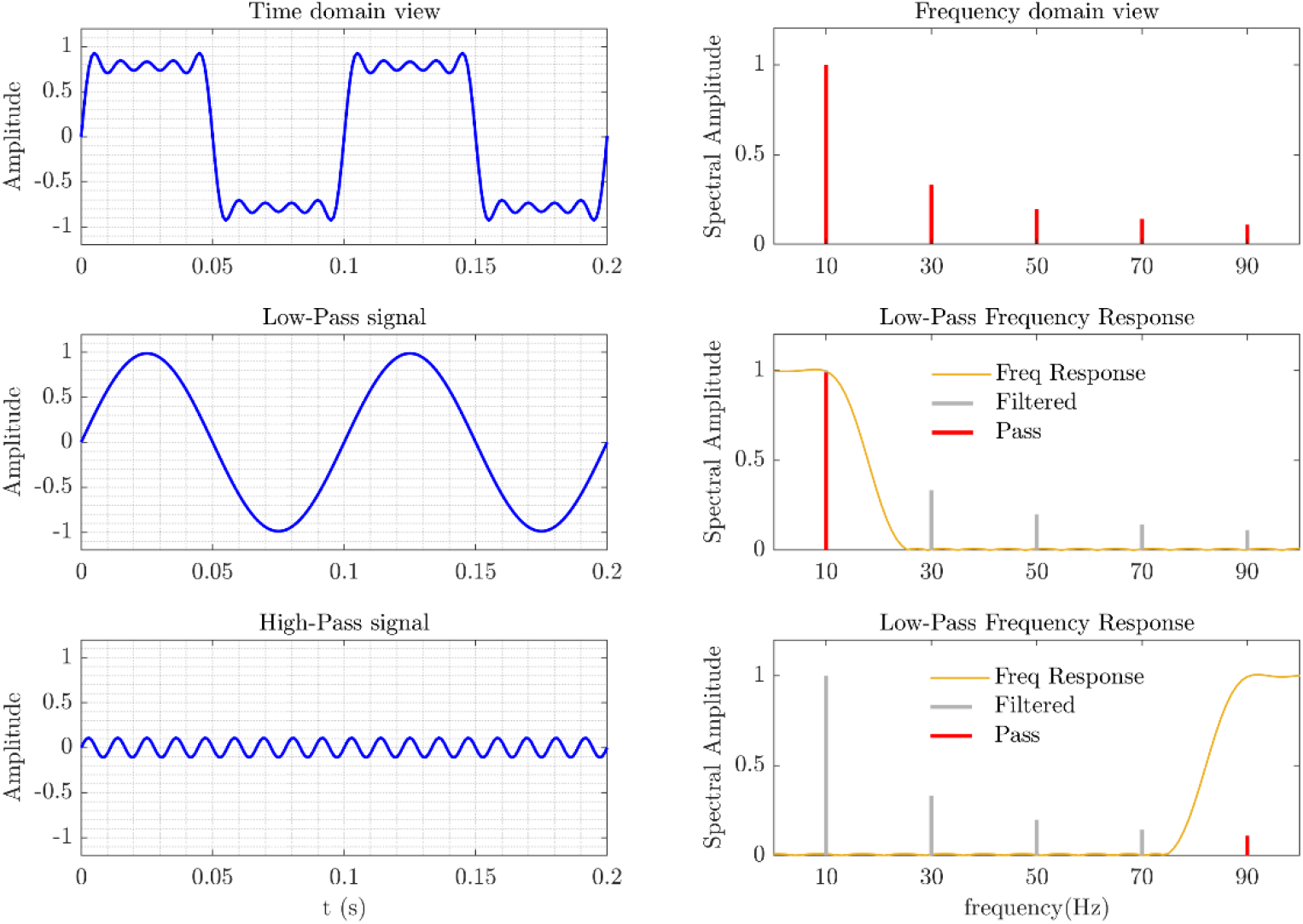
Exemplification of the filtering process. Top panel represents a signal composed by the sum of sinusoids and its spectral content. Middle panel illustrates the result of low-pass filtering, the frequency response allows the pass of spectral components below 10 Hz and attenuates spectral components above 25 Hz. The bottom panel shows the result of high-pass filtering, spectral components below 75 Hz are attenuated, since frequencies greater than or equal to 90 Hz are practically unchanged.

#### 2.3.2. Moving Average filters

Moving average filters can be considered as special cases of FIR filters, for that case the Eq. 5 represents the formulation of a moving average filter. If we compare Eq. 4 and Eq. 5, can be observed that filter coefficients *a_i_* = 1/*N* and *b*_*i*_ are zero. Moving average filters are simple for implementation and an effective choice for smoothing signals and for removing random noise, while maintaining signal trends [16][15].

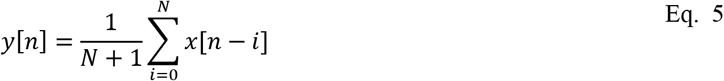

Figure 6 shows the application of moving average filter on sinusoidal signal containing random noise. After moving average filter, the resulting signal is similar to the original signal without noise, see Figure 6 B).

**Figure 6.**
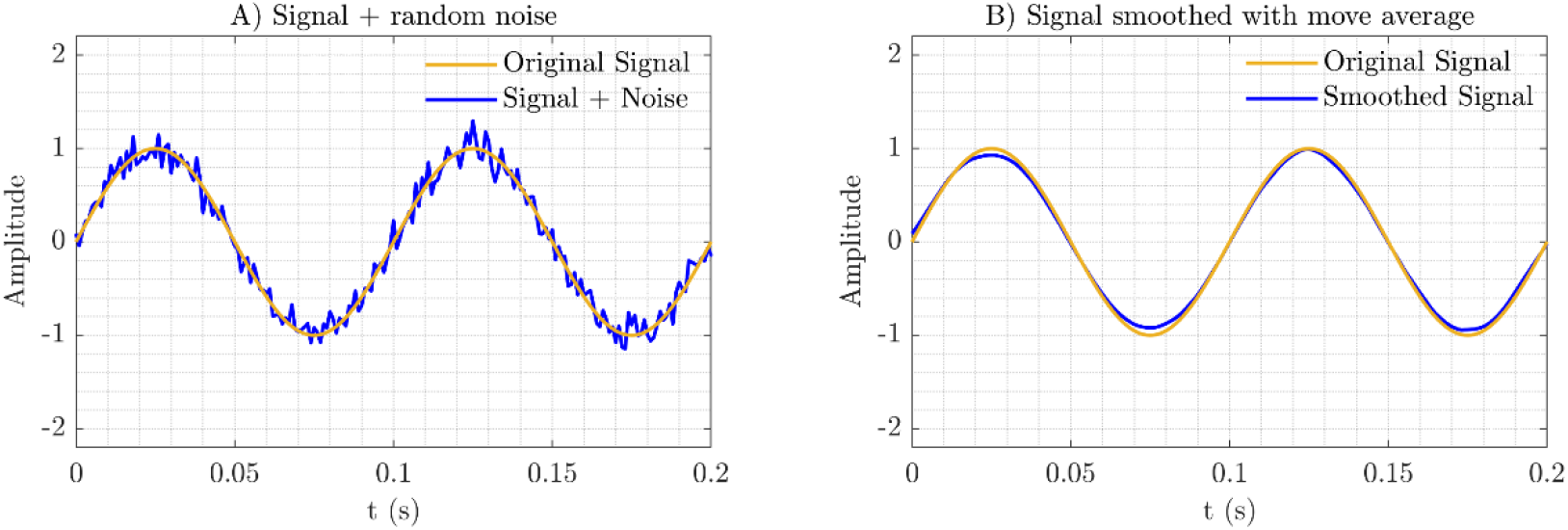
Application of moving average filter for smoothing a sinusoidal signal contained random noise. A) Noisy signal and original sinusoidal signal, noisy signal contains abrupt oscillations generated by random noise. B) Smoothed signal and original sinusoidal signal, smoothed signal after moving average filtering it is close to the original sinusoidal signal.

### 2.4. Parseval relation, Power Spectral Density and Sound Pressure Level

The Parseval’s relation states that the energy of a signal *x*[*n*] with N samples can be computed from the Fourier Transform. Specifically, if *X*[*k*] is the DFT of *x*[*n*], then the Parseval’s relation [6] can be written as in Eq. 6:

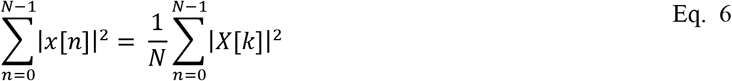

The Parseval’s relation is helpful in Ecoacoustics, since it can be used to compute energy variations in time signal through the frequency domain. Measurements related to signal energy variations could be used for acoustic events detection. Here we illustrate how to compute signal energy variation from frequency domain, but the main goal is not to detail the complete procedure to obtain calibrated measurements, for a more detailed explanation see [14].

The Mean Square Pressure (MSP) is related to the mean square value of a sound pressure signal *x*[*n*], and can be computed as in Eq. 7:

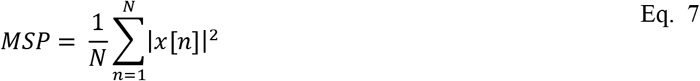

Power Spectral Density (PSD) is the contribution of power per unit of bandwidth [12], and the power in selected frequency bands can be used to estimated the contribution of specific spectral components. PSD can be computed as the magnitude square of DFT *X*[*k*] [17], thus it is derived from the Eq. 6 that we can compute energy from PSD based on Parsevaĺs relation.

Figure 7 exemplifies the use of Parseval relation to compute MSP from frequency domain, in that way we can compute broadband MSP or MSP in selected frequency bands. Figure 7 shows that computing MSP from time domain or frequency domain results in the same quantity.

**Figure 7.**
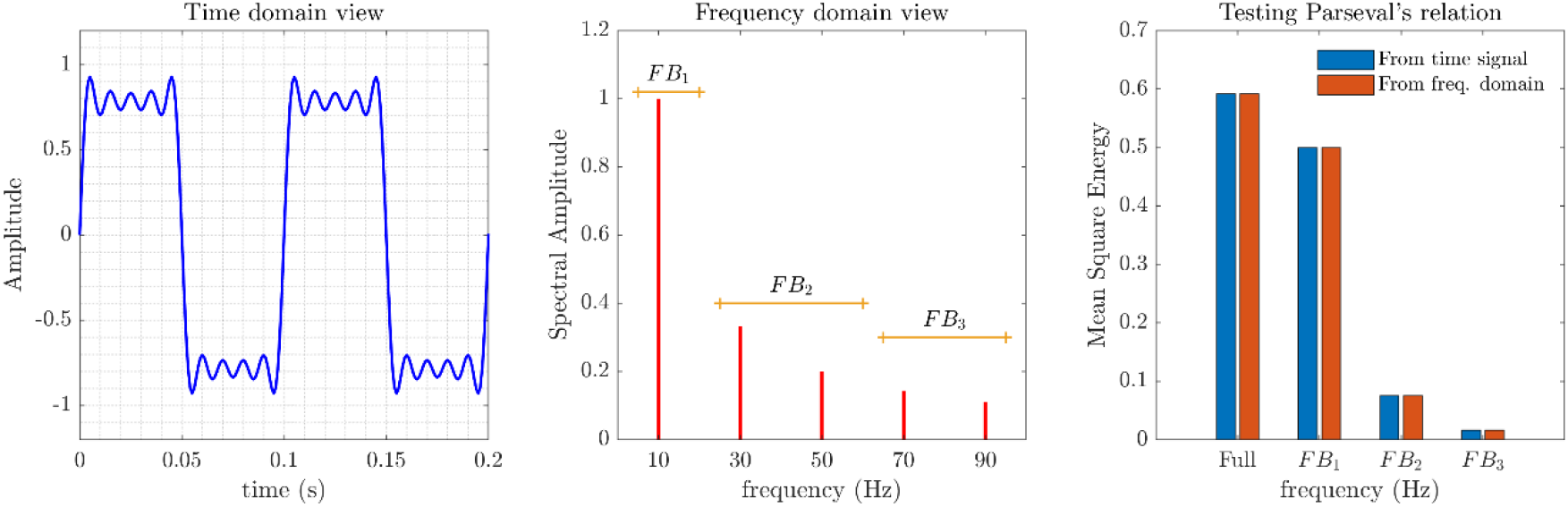
Testing the Parseval’s relation. A) Simulated signal composed by the sum of several sinusoidal waveforms. B) Spectral representation for the simulated signal, selected Frequency Bands (FB), named FB1, FB2 and FB3 are represented. C) The MSP is computed from the simulated signal (Full FB), and from the time signals representing the pure sinusoids in the respective frequency bands (FB1, FB2 and FB3). For example, to compute the MSP for the FB2: in time domain were used both the sinusoidal signals with 30 Hz and 50 Hz; from the frequency domain was summed the PSD values corresponding to spectral components comprised in FB2.

A useful application of Parsevaĺs relation is that we can compute energy or power in selected frequency bands, without the need to do the calculations with the signal in the time domain. Sound Pressure Level (SPL) is a common metric used in acoustics analyses [14], SPL can be expressed as in Eq. 8. This metric can also be computed from frequency domain, once calculated PSD matrices. The calculation and usefulness of PSD matrices will be illustrated in following sections.

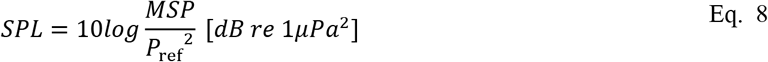

### 2.5. Pipeline for 24-h EDA

The procedure used to calculate daily PSD matrices, named here as Pxx matrices, are explained below and exemplified in Figure 8. For each wav file, recorded for the day j (j varying from February 4^th^ to March 4^th^) a respective pxx^†^ matrix was computed, by means of the Welch method [18], with 1-s Hamming window, 1025 frequency points, 50 % of overlap, and with 60-s temporal signal segments. Then, all pxx matrices referents to day j were merge in to a daily Pxx matrix and, finally, by visualizing Pxx matrices the 24-h spectrograms were obtained. Based on the system record settings, the numbers of recorded wav files by days, was N = 96. Aiming to analyses the whole collected dataset, 24-h spectrograms were constructed for each monitored day. These 24-h spectrograms were obtained by visualization of daily Pxx matrices.

**Figure 8.**
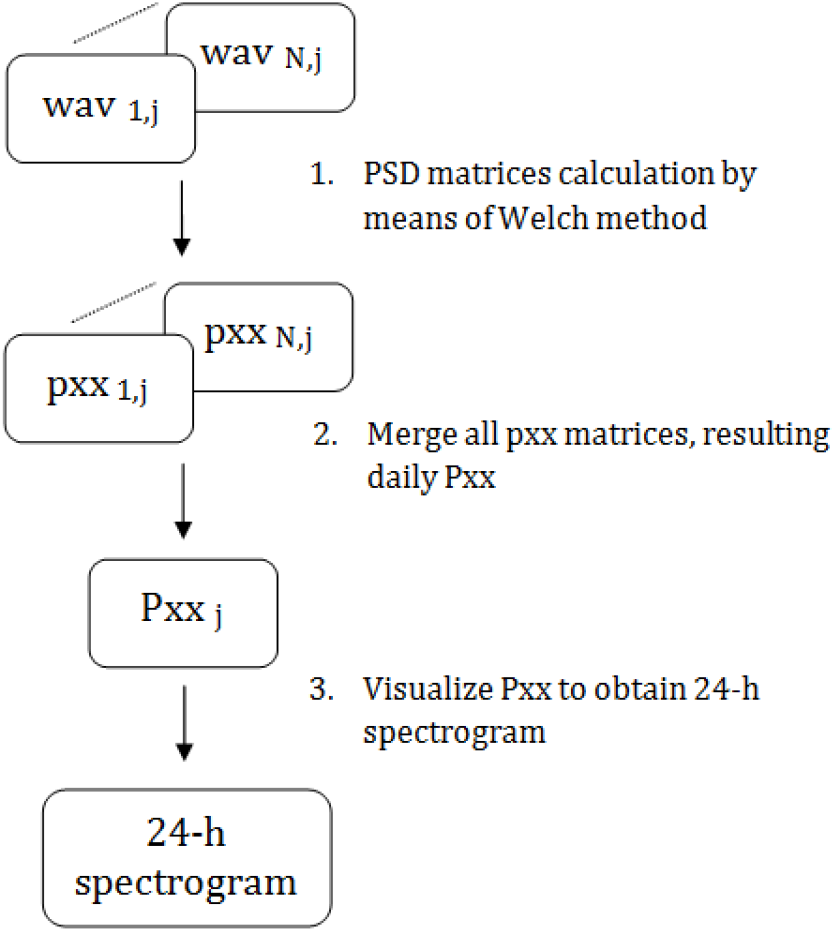
Flowchart for presented pipeline.

## 3. Some applications for 1-month underwater sound data

Here we describe as a case study 1 month of continuous underwater sound recorded at Xixová-Japuí State Park (XJSP), located on the SW of Santos Estuarine System (São Paulo State, Brazil), see Figure 9. This area has a rich daily soundscape, but it is also affected by urbanization, industrial and port activities [19]. Fish choruses with daily periodicities were the most important contributor for the XJSP soundscape [19], [20]. The location for the monitored place is represented in Figure 9. Underwater acoustic data was recorded by means of an autonomous passive monitoring system [21]. The acoustic signals were acquired at 11.025 kHz sample rate, with 16-bit resolution; individual sound files of 15 min duration were continuously stored in a SD card.

**Figure 9.**
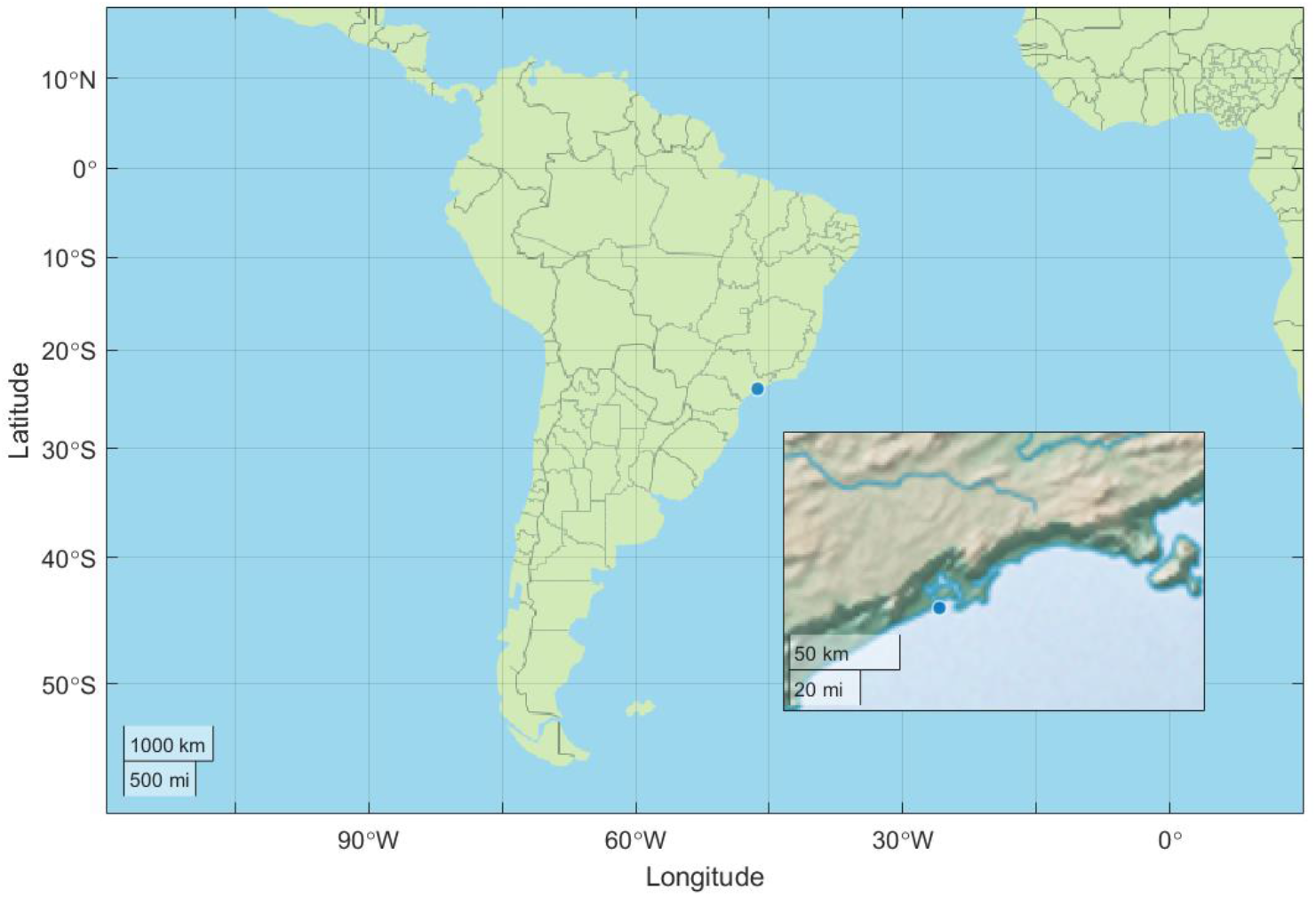
Monitored point located at Xixova marine park.

### 3.1. 24-h Spectrogram and 24-h SPL

24-h spectrograms are valuable for visualizing daily soundscape variations for a given monitored area. Averaging 24-h spectrogram also can be used for summarizing soundscapes [19][22] for representative periods, as for example 1 month. Dily 24-h spectrograms for complete four weeks are illustrated in Figure 10. It is worth nothing that acoustic events seem to occur every day with a precise time-spectral patters. For summarizing sound data recorded for 1 month was computed the mean 24-h Spectrogram, illustrated in Figure 11. From 24-h mean spectrogram, four main acoustic events can be noticed, these acoustic events are originated by fish choruses and are denoted as Ch_1_, Ch_2_, Ch_3_ and Ch_4_.

**Figure 10.**
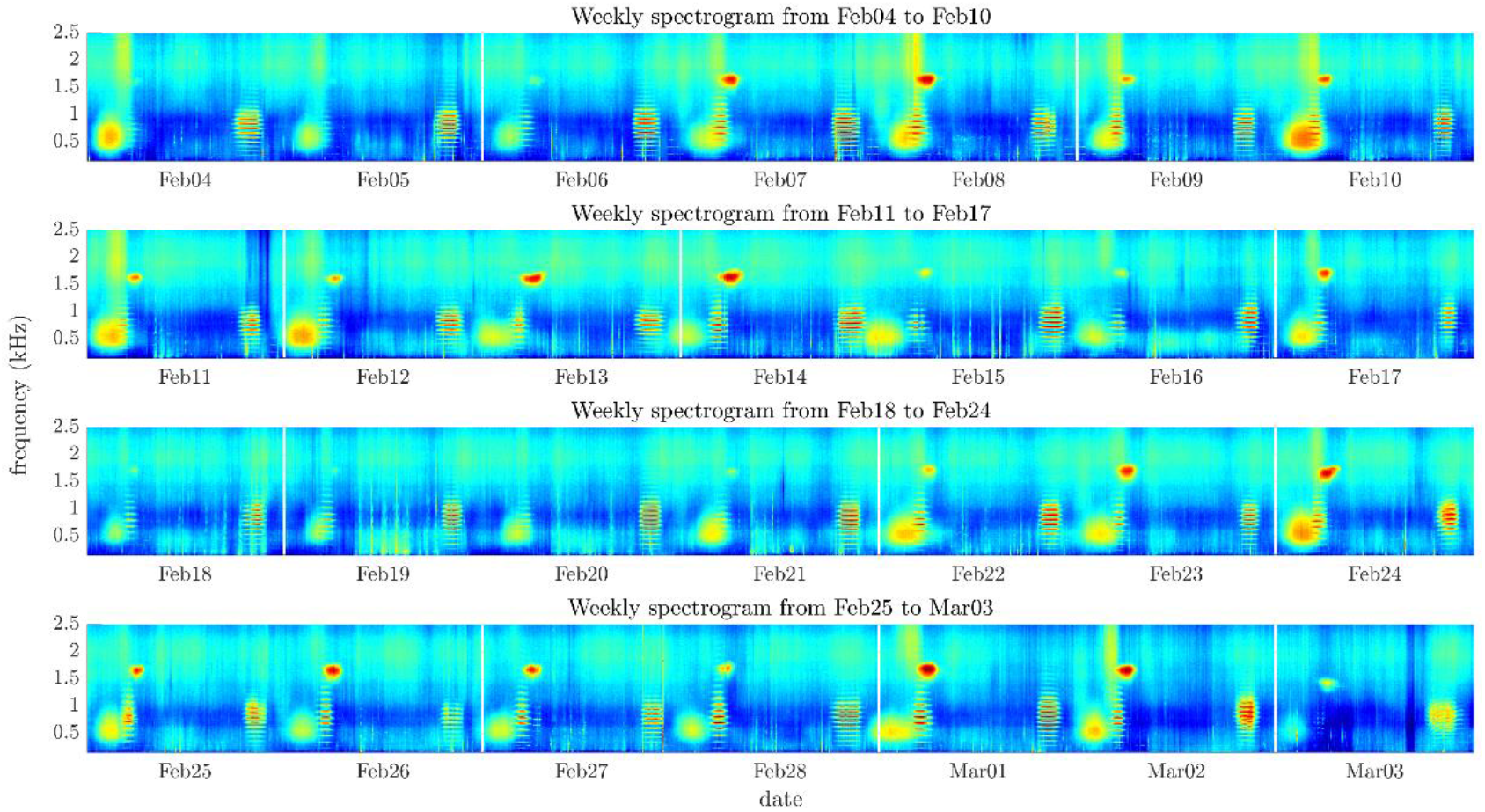
Four weeks spectrograms. The weekly spectrograms allow to identify acoustic events that suggest a daily pattern.

**Figure 11.**
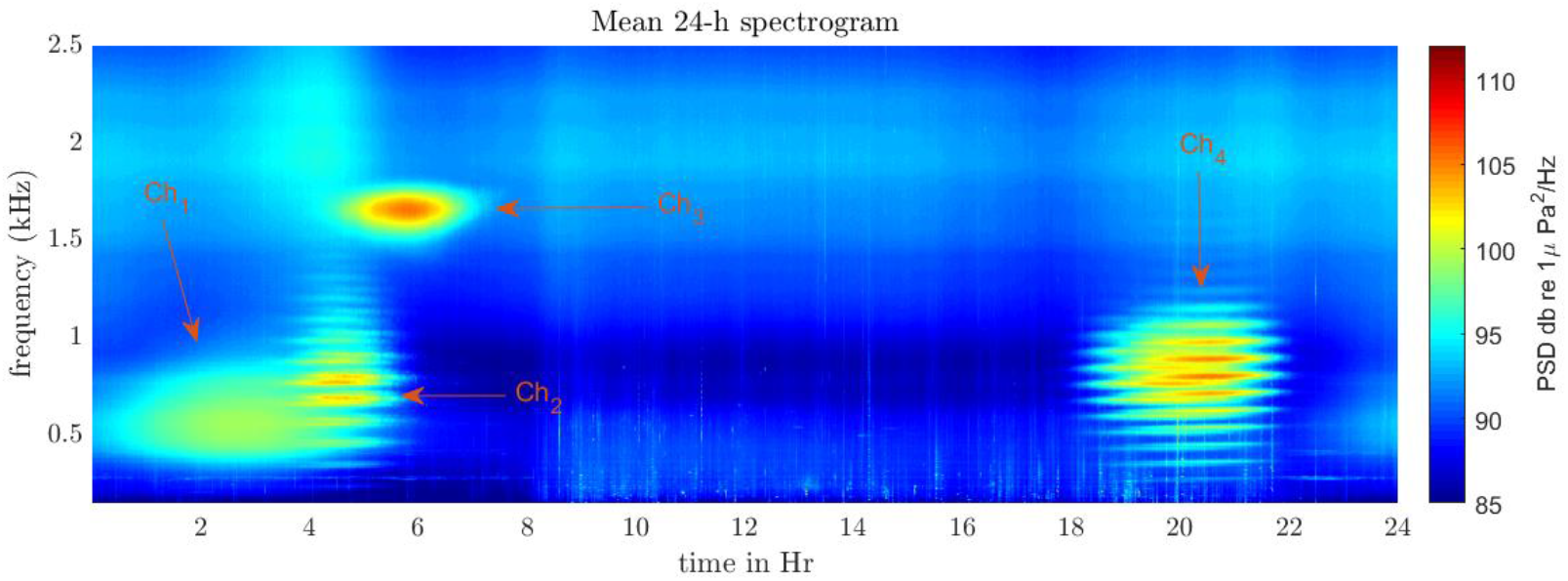
Mean average for 24-h spectrograms between February 04 to March 04.

SPL computed within the Low-Frequency Band (Low-FB) and High-Frequency Band (Hig-FB) were also used for studying sound levels variations. For the analysis was selected a Low-FB that comprises frequencies form 150 Hz to 1.2 kHz and Hig-FB that includes frequencies from 1.5 kHz to 2.0 kHz. Figure 12 shows four-week Low-FB SPL time series and Figure 13 represent four-week Hig-FB SPL time series.

**Figure 12.**
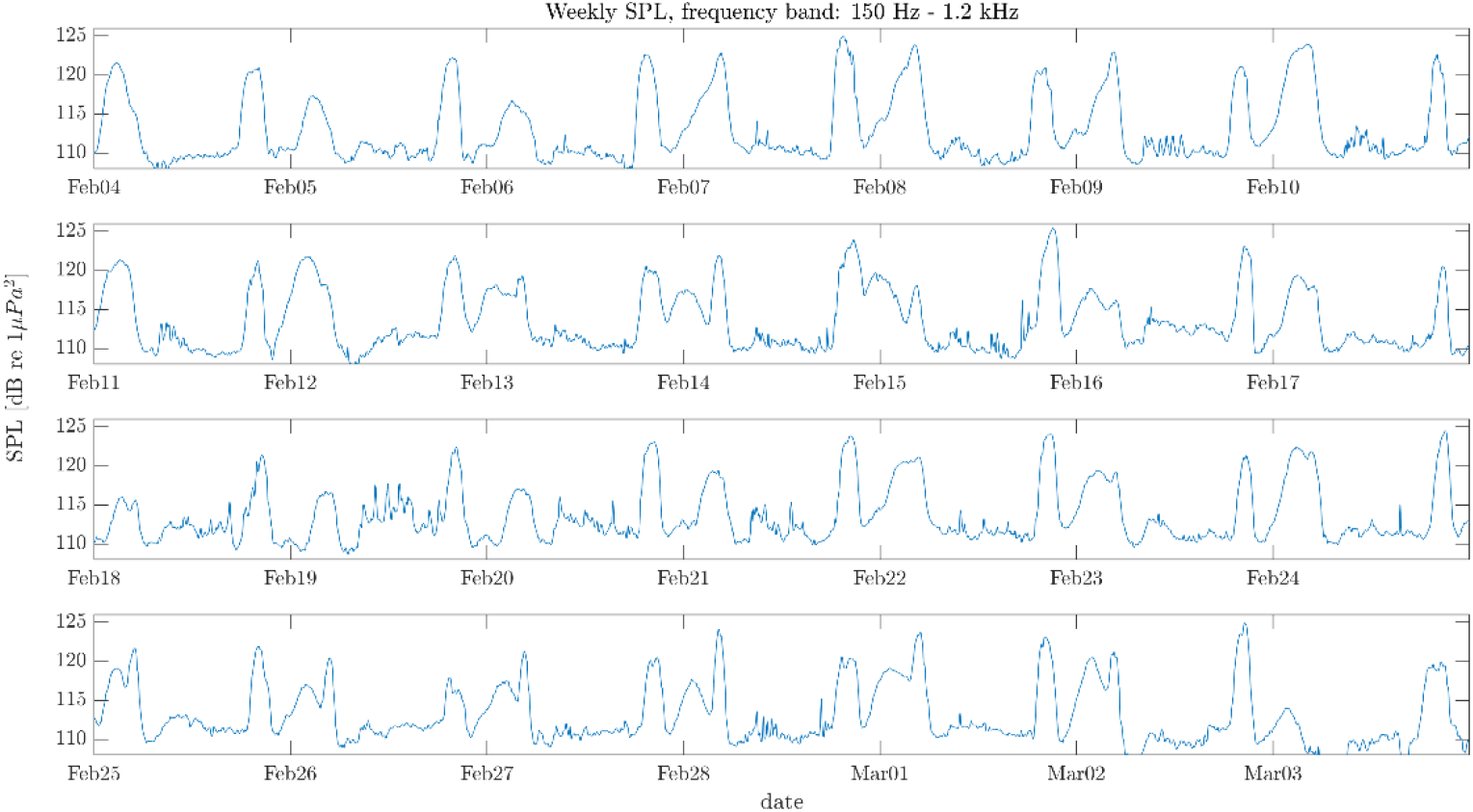
For week Low-FB SPL, each panel represents one week, respectively.

**Figure 13.**
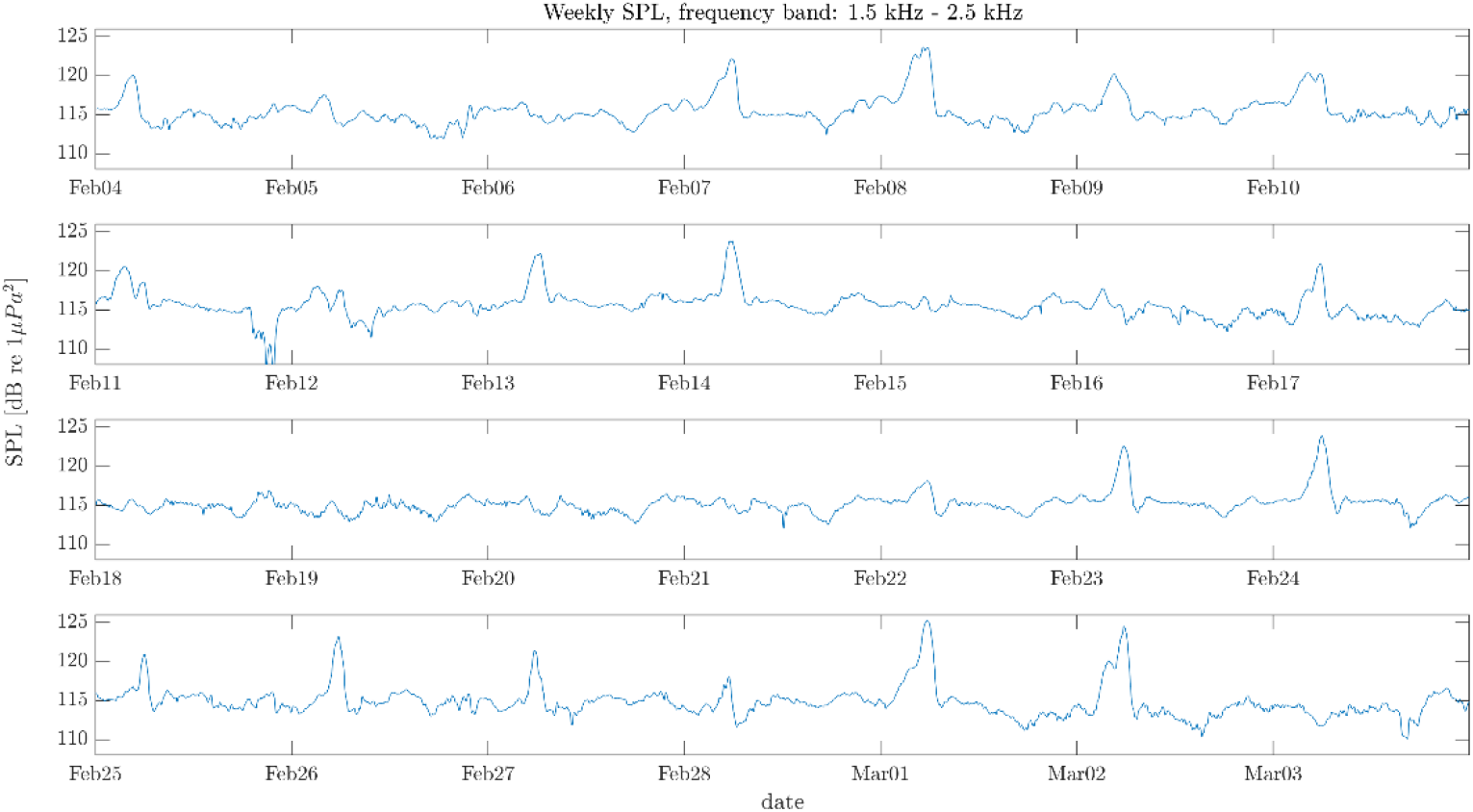
For week Hig-FB SPL, each panel represents one week, respectively.

### 3.2. Finding soundscape periodicities

The auto-covariance and the spectral estimation of the SPL time series were used for quantifying perceptible periodicities in the analyzed data. Figure 14 shows the auto-covariance and the spectrum Low-FB SPL and Hig-FB SPL, respectively. For the purposes of this work were defined strong periodicities and weak periodicities as principal peak and secondary peaks detected on PSD curve. The strong periodicities are annotated in red and weak ones are annotated with green markers, both in time domain (auto-covariance) and in frequency domain (spectral content illustrated trough the PSD). For estimating periodicity values trough time domain analysis, was determined the mean inter-interval between peaks (in minutes). By computing the ratio between the number of minutes for one day (1440 = 24h*60min) and the mean inter-interval was finally estimated the periodicities implicit in SPL time series. On the other hand, in frequency domain finding periodicities can be done in a simple way, by detecting the prominent spectral peaks.

**Figure 14.**
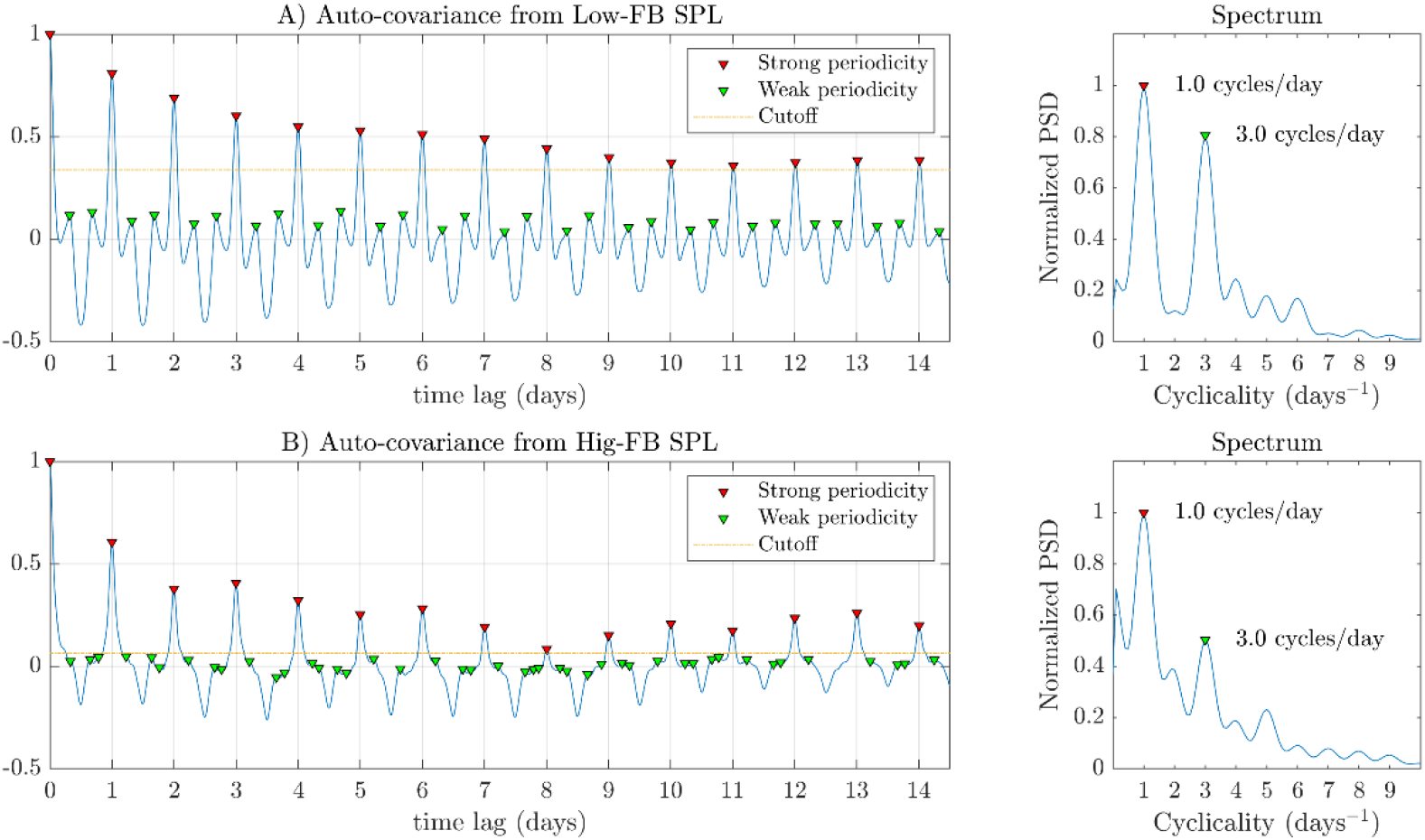
Auto-covariance and the spectrum of Low-FB SPL and High-FB SPL. Peaks in red markers indicates strong periodicities and green markers point out week periodicities. A cut-off line indicates the minimum value for strong periodicity peak in auto-covariance.

For the case of Low-FB SPL both the strong periodicities and weak periodicities computed from time domain (auto-covariance) or from frequency domain (by means of PSD) results in almost the same value, 1 cycle/day and 3 cycles/day, respectively. Thus, for Low-FB SPL the computed strong periodicity is related to a daily cycle, which agrees with the repetitive 24-h patterns that can be visually detected in Figure 10. For the case of weak periodicity detected as 3 cycles/day, we can see from mean 24-h spectrogram (Figure 11) that in Low-FB (150 Hz – 2.0 kHz) there are 3 different major acoustic events, which can explain the occurrence of that weak periodicity.

For the case of Hig-FB SPL the strong periodicity computed from time domain or spectral estimation is the same, but this is not the case for computed weak periodicity. The weak periodicity computed from time domain was approximately 4.1 cycles/day, which does not agree with the value computed from frequency domain (see Figure 14, bottom panels). The weak periodicity quantified for Hig-FB SPL could be related to weak events in this frequency band or also by the incidence of remaining spectral components for the 3 major events quantified in Low-FB.

### 3.3. SPL-gram

The SPL-gram representation allows to visualize the SPL variations for long-term acoustic monitoring [23]. By using the SPL-gram for selected frequency bands a "picture" of the temporal structure of the soundscape can be obtained. To obtain the SPL-gram, daily SPL for specific frequency band is smoothed, thus removing transient oscillations as for example ship noise, while preserving lasting events, such as fish choruses. Once daily smoothed SPL are obtained, the time series are converted to color bars. Finally, by merging color bars in chronological order the SPL-gram representation is obtained. Figure 15 shows the procedure for obtaining SPL color bars for the case of Low-FB SPL and Hig-FB SPL for a specific day.

**Figure 15.**
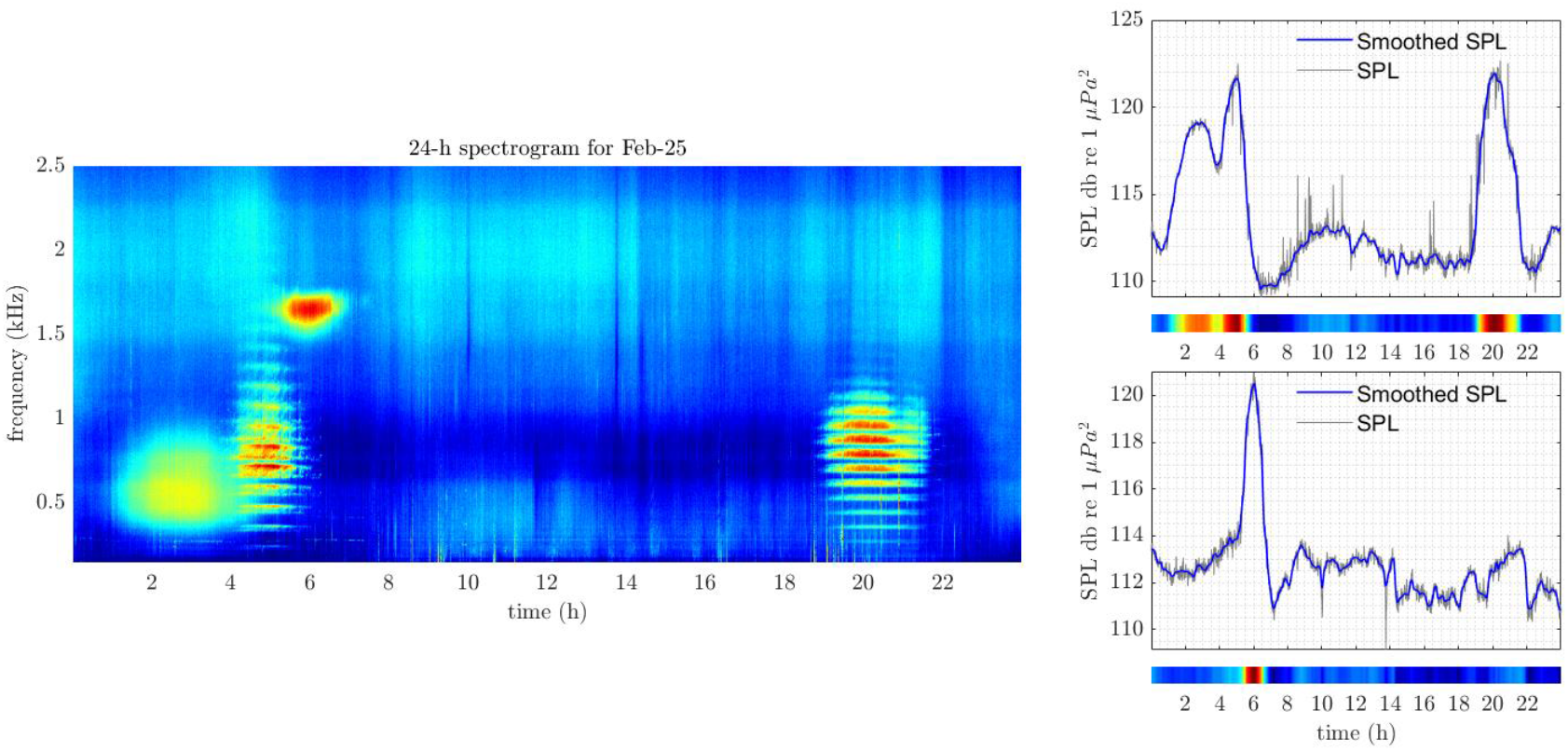
SPL-<u>gram</u> construction. Each 24-h SPL is converted to colorbar, then colorbars for each consecutive day are chronologically stacked. Left panel shows the spectrogram for February 25. Right panels represent Low-FB SPL (top panel) and High-FB SPL (botton panel).

Figure 16 shows the SPL-gram for Low-FB SPL and Hig-FB SPL. By plotting on the graph events occurrence such as sunrise and sunset we can obtain a temporal relation between soundscape variation and these events. For example, for the case of Low-FB SPL-gram, the occurrence of sound level increase around the time interval from 19 h – 21 h (UTC time) ends before sunset. This fact is related to the occurrence of Ch_4_ (see annotation in Figure 11), which seems to decrease its activity shortly before sunset. Another soundscape trend that can be observed for Low-FB SPL is that the period between 06 h – 18 h appears without substantial lasting acoustic events.

**Figure 16.**
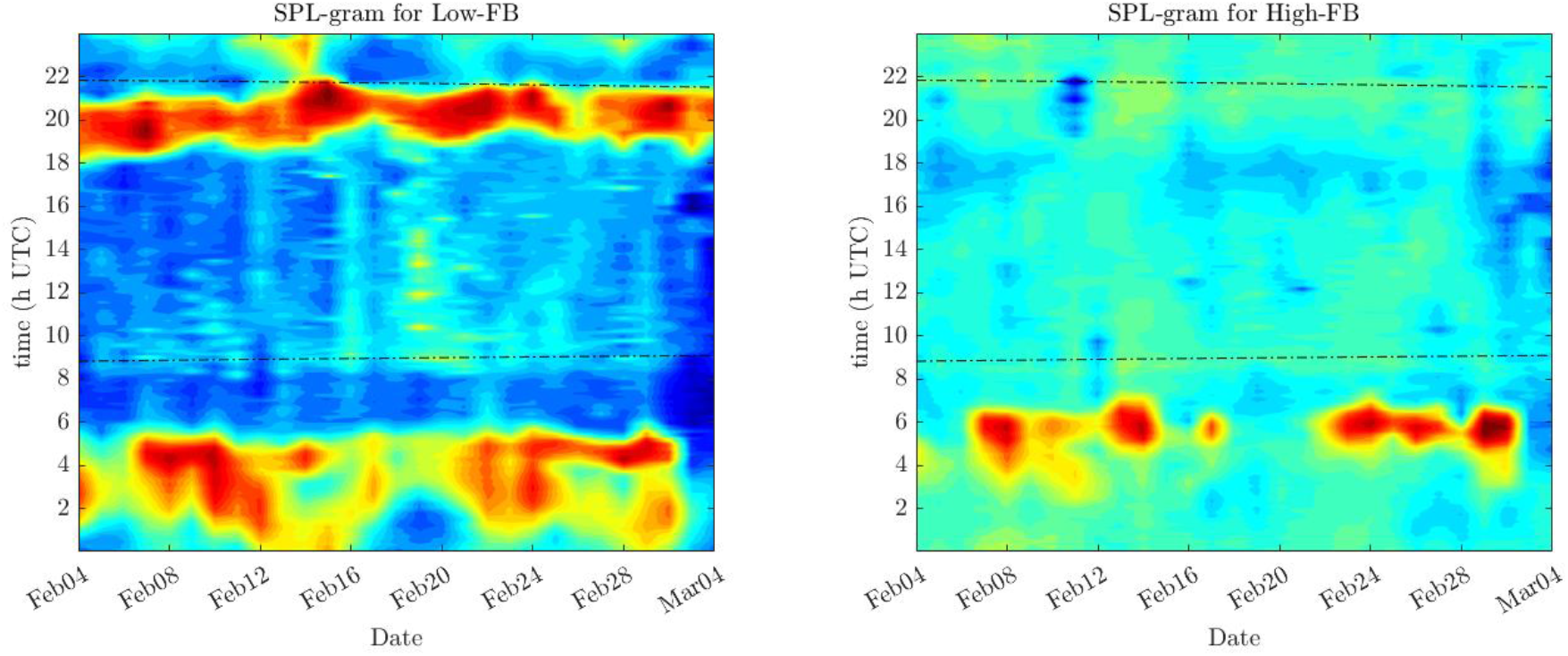
SPL-gram for continuous underwater sound recordings collected in Xixova State park, black dashed lines represent sunrise and sunset.

Also, mean values for daily SPL could be used for summarizing daily trends in the studied place. Figure 17 represents the mean of SPL time series, both in conventional plot (top panels) as in polar form (bottom panels). For Low-FB SPL can be observed two significant deviations from the baseline levels. Between 02 h - 05 h the first increase in Low-FB is noted, which in turn can be divided into two different events, one achieving the acoustic peak between 02 h – 03 h, and the other attaining the maximum level between 04 h - 05 h. The later deviation in Low-FB SPL it is reached between 20 h – 21 h. For the case of Hig-FB SPL one major deviation from baseline levels appears between 05 h – 06 h. These acoustic peaks agree with the visual information illustrated in mean 24-h spectrogram and the events annotated as Ch_1_, Ch_2_, Ch_3_ and Ch_4_ (Figure 11).

**Figure 17.**
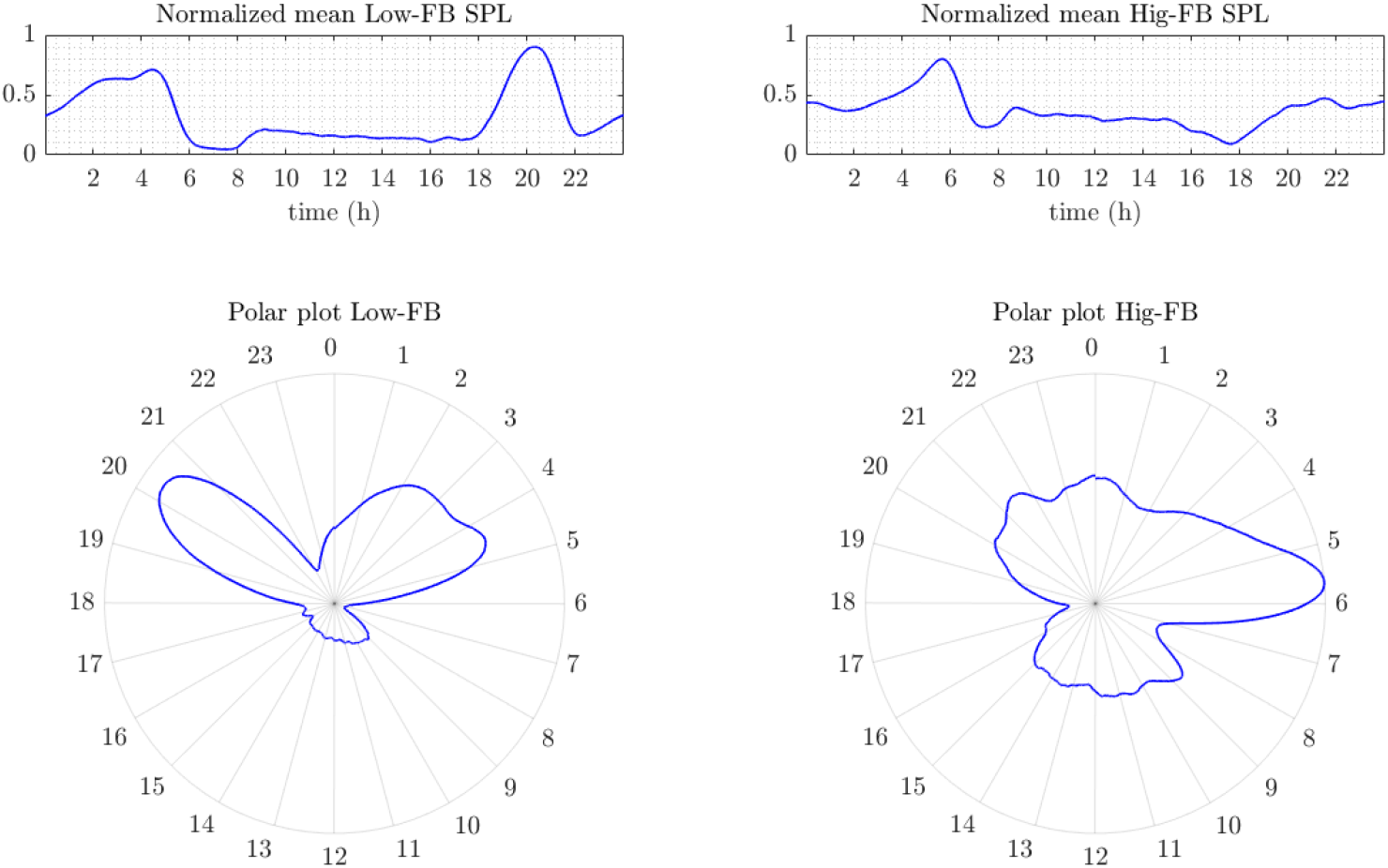
Polar plot representation for normalized mean Low-FB SPL and Hig-FB SPL. Top panels represent the normalized mean SPL, both for Low-FB and Hig-FB in the conventional amplitude vs time plot. Bottom panels illustrate the same information as top panels, but in polar plot.

## 4. Conclusion

Results showed that basic concepts of the field of digital signal processing are valuable tools for Ecoacoustic researchers. To name a few, understanding basic concepts such as filtering, spectral analysis, and visualizations such as spectrogram can be particularly useful for explorative data analysis in Ecoacoustics. Thus, the comprehension of those concepts and the knowledge about how to use it in real applications could contribute to the field.

The concepts presented in the present manuscript are explained in more detail in technical books and papers [6][7]. However, the main objective of this work was to present the concepts in a comprehensible way, be means of visualizations and straightforward explanations focused for the specific area of Ecoacoustics. The concepts presented were used for application on real sound data, composed by 1-month underwater recordings. Here was demonstrated the utility of tools such as 24-h spectrograms and mean 24-h spectrogram por visualizing daily variations and for summarizing soundscape patterns. Also, the use of metrics such as SPL results useful for quantifying periodicities that could be visually perceived on spectrogram representations. The SPL-gram evidenced to be useful as soundscape visualization tool, allowing a graphical representation for acoustic patterns summarization. Finally, normalized SPL an its representation in polar form showed to complement the comprehension and insights obtained from 24-h mean spectrogram and SPL-gram.

## Acknowledgments

Author would like thank Linilson Padovese for discussions and major contributions that allowed the development of the current manuscript.

## Footnotes

* Hamming function (as well as Hanning, Bartlett and others possible windows) has values close to zero at the extremes and close to one at the middle. This justifies why the multiplication by the window function diminishes possible discontinuities between beginning and ending of the windowed signals.

† pxx matrix refer to PSD matrix computed for individual sound files, while Pxx matrix refers to daily period, thus Pxx for specific day is obtained by merging all pxx matrices for that day.

## Notes

### Competing Interest Statement

The authors have declared no competing interest.

### Summary of Updates

Fixed some typos and updated some figures

